# Dopamine promotes *Klebsiella quasivariicola* proliferation and inflammatory response in the presence of macrophages

**DOI:** 10.1101/2023.01.28.526064

**Authors:** Xiang Li, Lin Cheng, Xueyang Liu, Xiaoli Wang, Rui Li, Shao Fan, Qiulong Yan, Tonghui Ma, Yufang Ma, Jian Kang

## Abstract

*Klebsiella quasivariicola* was a novel strain of *Klebsiella* species and had potential pathogenicity. Our previously studies showed dopamine, one of the most commonly used rescue drugs for critically ill patients, had clear effects on the growth of *K. quasivariicola* in culture medium, however, its effects on host immune system were ignored. Therefore, in consideration of the host immunity, the interactions of *K. quasivariicola*, dopamine and macrophages were explored. In this study, RAW264.7 cells and C57/BL6 mice were infected with *K. quasivariicola*, and the bacterial growth in macrophage, the production of inflammatory cytokines and the pathological changes of mice lung were detected, in the absence or presence of dopamine. Our results showed dopamine inhibited the *K. quasivariicola* growth in medium, but promoted the bacterial growth when *K. quasivariicola* was co-cultured with macrophages; The expression of proinflammatory cytokines in *K. quasivariicola* infected RAW 264.7 were increased, while a sharp rise was observed with the addition of dopamine; Infection of *K. quasivariicola* to mice induced an inflammatory response and lung injury, which were exacerbated by dopamine administration. It can be concluded that dopamine administration resulted in a significantly increase of *K. quasivariicola* burdens in the presence of macrophage, consequently, aggravated the inflammatory response and inflammatory injury.

**Importance:** Dopamine is one of the most commonly used rescue drugs for critically ill patients. Here we indicated *K. quasivariicola* was a potential pathogen of pulmonary infection, and dopamine significantly increased the proliferation of *K. quasivariicola* when exposed to macrophage, subsequently result in severe inflammatory response and inflammatory injury. We also proposed an *in vitro* model of microbes-drugs-host immune cells that could better mimic *in vivo* environment and more suitable for the studies of inhibitor screening. This fundamental work had contributed to the present understanding of the crosstalk between pathogen, dopamine and host immune cells. Furthermore, our data showed dopamine was one of the risk factors for patients with *K. quasivariicola* infection, which provided a basis for clinical precision medicine.

## Introduction

Nosocomial infection is a serious problem in intensive care units (ICU). It can cause the increase hospitalization time, risk of complications, and also the additional financial burdens on patients (1). The risk factors of nosocomial infection include the use of invasive treatment, the administration of hormones and antibiotics (2, 3). In addition, recently, examples have been reported that dopamine, a commonly used drug in ICU, was also associated with infections. Dopamine (DA), one of the most potent catecholamine vasopressors, is widely used in various clinical settings, especially in antishock therapy to critically ill patients. However, recently, several clinical studies indicated the relationship between dopamine and infections, made it necessary to exercise caution when using dopamine (4-6). A randomized controlled trial of pediatric septic shock showed the patients treated with dopamine showed a higher rate of infections than in those treated with epinephrine (7). Similar, Hatachi *et al*. (8) reported that the administration of dopamine was a risk factor for infection in children after cardiac surgery. They indicated that the nosocomial infection rates were related to both the duration of dopamine administration and the total dopamine dose. In addition, a retrospective study of extremely preterm infants demonstrated that the increased amount of dopamine was associated with infection (9). Furthermore, i*n vivo* and *in vitro* tests also confirmed that dopamine could affect the growth or virulence of certain bacteria (10, 11). All these indicated dopamine treatment was linked to infection, in an ambiguous way.

Culturomics has facilitated in the large-scale isolation of human bacterial strains (12) and has provided the foundation for bacterium-drug interaction research (13, 14). Recently in our lab, based on the culturomic methods, a group of potential pathogenic bacteria were isolated from sputum sample of the pneumonia patients from ICU. To explore the relationships between dopamine and these potential pathogenic bacteria, the growth of some isolated bacteria was detected with the presence of dopamine (Fig S1). Among them, the growth of *Klebsiella quasivariicola*, a novel strain of *Klebsiella* species, was clearly impacted by dopamine, which engrossed our attention (Fig S1).

*Klebsiella quasivariicola* was recently sequenced from a human clinical isolate (15). The whole-genome sequencing data showed that there was an extended-spectrum β-lactamases enzyme gene in *K. quasivariicola* genome, which suggested its potential for causing the serious human infections (15). Later, *K. quasivariicola* strains were successively isolated from the wound infections of a diabetic foot infected patient and the urine samples of community-acquired infections (16, 17). Our results indicated that *Klebsiella quasivariicola* was a potential pathogen in pulmonary infection. Other studies also confirmed that *K. quasivariicola* strain was multidrug resistant, which was resistant against norfloxacin, ciprofloxacin, cefazolin and vancomycin (16), indicating that *K. quasivariicola* was an important potential pathogen and could not be underestimated. However, the exact mechanism by which *K. quasivariicola* infection remains largely unknown.

Our previously study has clearly showed that dopamine could affect the growth of *K. quasivariicola in vitro* (Fig S1A). However, during the interactions between dopamine and bacteria, one of the important factors, the host immune system, might be ignored. In host, alveolar macrophage is the first sentinel to defense against pathogen in lung. Although the numbers of alveolar macrophages are much less in alveoli, these cells can constantly patrol among multiple alveoli and eradicate pathogen independent of neutrophil recruitment (18). Alveolar macrophages can effectively avoid the inappropriate inflammation and injury and also maintain the pulmonary homeostasis, however, when large amount or highly virulent pathogens are exposed, a potent inflammatory response will be triggered. The overstimulated macrophage will release a variety of proinflammatory cytokines such as IL-6 and TNF-α, creating a cytokine storm which cause tissue injury, pulmonary dysfunction or even death (19). In clinic, especially in ICU, the infection of *Klebsiella app*. always leads to a robust inflammatory response, which brings a great threat to patients (20). Therefore, it is wondered that if *K. quasivariicola* could provoke a macrophage inflammatory response, and if dopamine has any effect on this process.

In this study, the effects of dopamine on the growth and potential pathogenicity of *Klebsiella quasivariicola* was investigated by using both macrophage cell line of RAW264.7 and mice model. Our results showed dopamine inhibited the growth of *K. quasivariicola* in culture medium but promoted the viability of *K. quasivariicola* in the precence of macrophages. The *K. quasivariicola* strain could lead to pulmonary inflammation. Significantly, the administration of dopamine elevated the expression of pro-inflammatory factors, thereby exacerbated *K. quasivariicola* infection. This fundamental work had contributed to the present understanding of the crosstalk between pathogen, dopamine and host immune cells. Furthermore, our data showed dopamine was one of the risk factors for patients with *K. quasivariicola* infection, which provided a basis for clinical treatment.

## Results

### Dopamine promoted the proliferation of *K. quasivariicola* when bacteria were co-cultured with macrophages

To determine the effects of dopamine on *K. quasivariicola* growth, the bacterial growth was measured in the presence of dopamine. When cultured in DMEM Basic medium, the growth of *K. quasivariicola* treated with 500 μg/ml dopamine exhibited a certain degree of decline compared to that of bacterial culture without dopamine (Fig.1A,B). But after incubation with RAW264.7 for 4 h (MOI=10:1), the *K. quasivariicola* numbers had a significant increase with the supplement of 500 μg/ml dopamine, both in cell culture medium (Fig. 1C) and inside the macrophages (Fig. 1D). These indicated dopamine could promote the proliferation of *K. quasivariicola* in the presence of macrophages.

**Figure 1:**
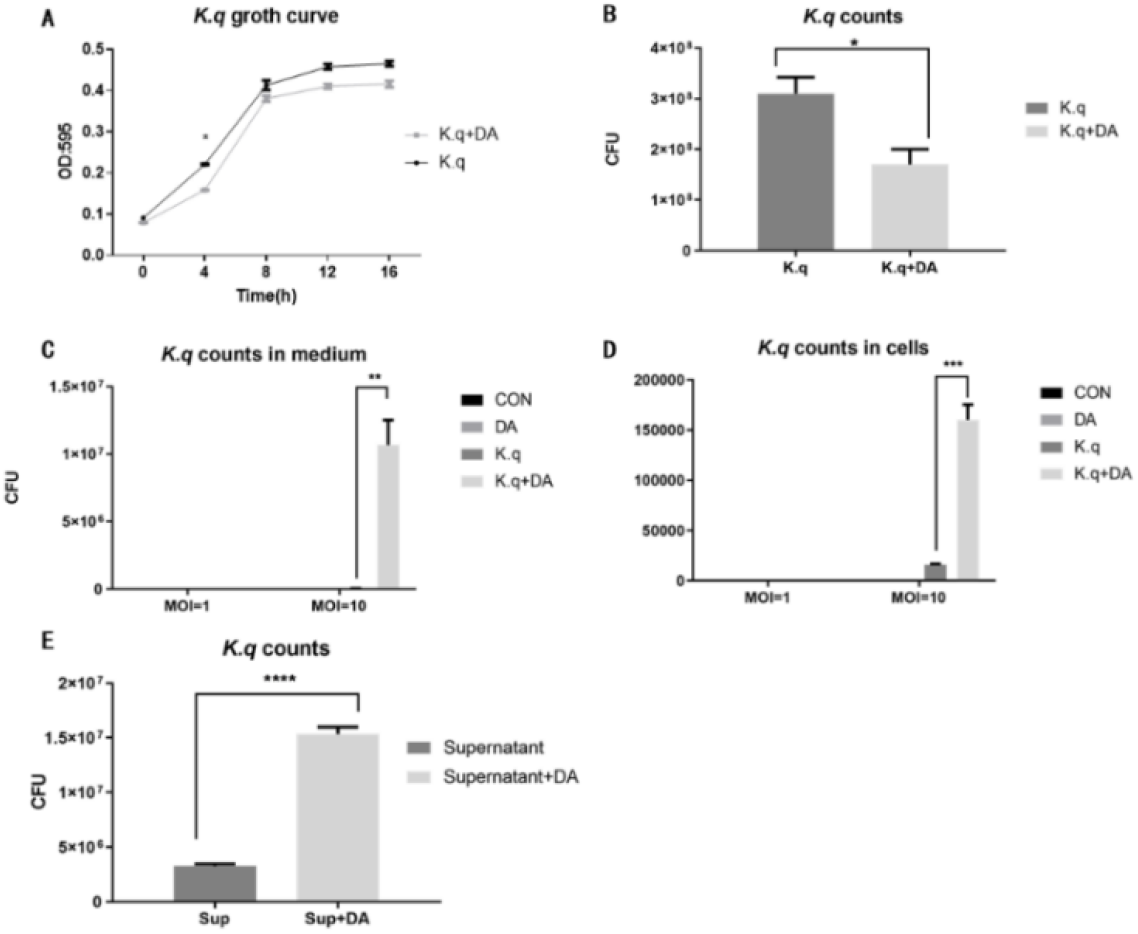
Growth curve and CFU count of *K. quasivariicola* under different conditions. (A-B) The growth curve (A) and CFU counting (B) of *K. quasivariicola* in DMEM Basic medium. Black, *K. quasivariicola* cultured only; Gray, *K. quasivariicola* cultured with 500 µg/ml dopamine. (C-D). After infecting RAW 264.7 cell with *K. quasivariicola*, CFU counts of *K. quasivariicola* in cell culture medium (C) and in macrophages (D) were performed. (E) CFU of *K. quasivariicola* grown in supernatant of RAW 264.7 culture medium. RAW264.7 cells were cultured in DMEM medium containing 10% FBS, in the absence (black) or presence (gray) of 500 μg/ml dopamine. The supernatants of cell culture medium were collected and used to cultivate *K. quasivariicola*. The CFU were determined after 8 hours of cultivation. The OD620 of cultures were monitored every 4 hours. Each sample was assayed in triplicate. Asterisk (*) means the difference between two groups was statistically significant.

Moreover, the supernatants produced by culturing RAW 264.7 were also collected for *K. quasivariicola* cultivation. RAW 264.7 cells were cultured in DMEM Basic medium with 500 μg/ml dopamine for 4 hours, and then the culture supernatants (cell free) were collected. Interestingly, the supernatants of RAW 264.7 cells cultured in the presence of dopamine showed a great capacity to enhance the growth of *K. quasivariicola* (Fig.1E).

### Dopamine significantly elevated the levels of pro-inflammatory factors in RAW 264.7 in response to *K. quasivariicola* infection

To detected the inflammatory responses triggered by *K. quasivariicola*, a series of cytokines produced by RAW 264.7 were measured after *K. quasivariicola* infections. Real-time PCR showed the mRNA levels of TNF-α, IL-6 and IFN-γ in RAW 264.7 of *K. quasivariicola* group were higher than control group after infection for 4 and 12 hours, respectively (Fig. 2A). After 4 hours-infection with *K. quasivariicola*, the mRNA levels of iNOS, IL-6, TNF-α, IFN-γ, chemokines CXCL1, CXCL2, and the NLRP3 inflammasome were all increased in RAW 264.7 cells (Fig. 2B). It was worth noting that the mRNA levels of these pro-inflammatory cytokines were significantly elevated when dopamine were added (Kq+DA group) (Fig. 2). Moreover, immunofluorescence co-localization analysis revealed that NLRP3 expression was obviously increased when RAW 264.7 was infected with *K. quasivariicola* (K.q), and that NLRP3 expression was further risen when dopamine was added (K.q+DA) (Fig. 2C). All these indicated that *K. quasivariicola* could trigger the production of a series of pro-inflammatory factors in RAW 264.7, and the use of dopamine would exacerbate the inflammation.

**Figure 2:**
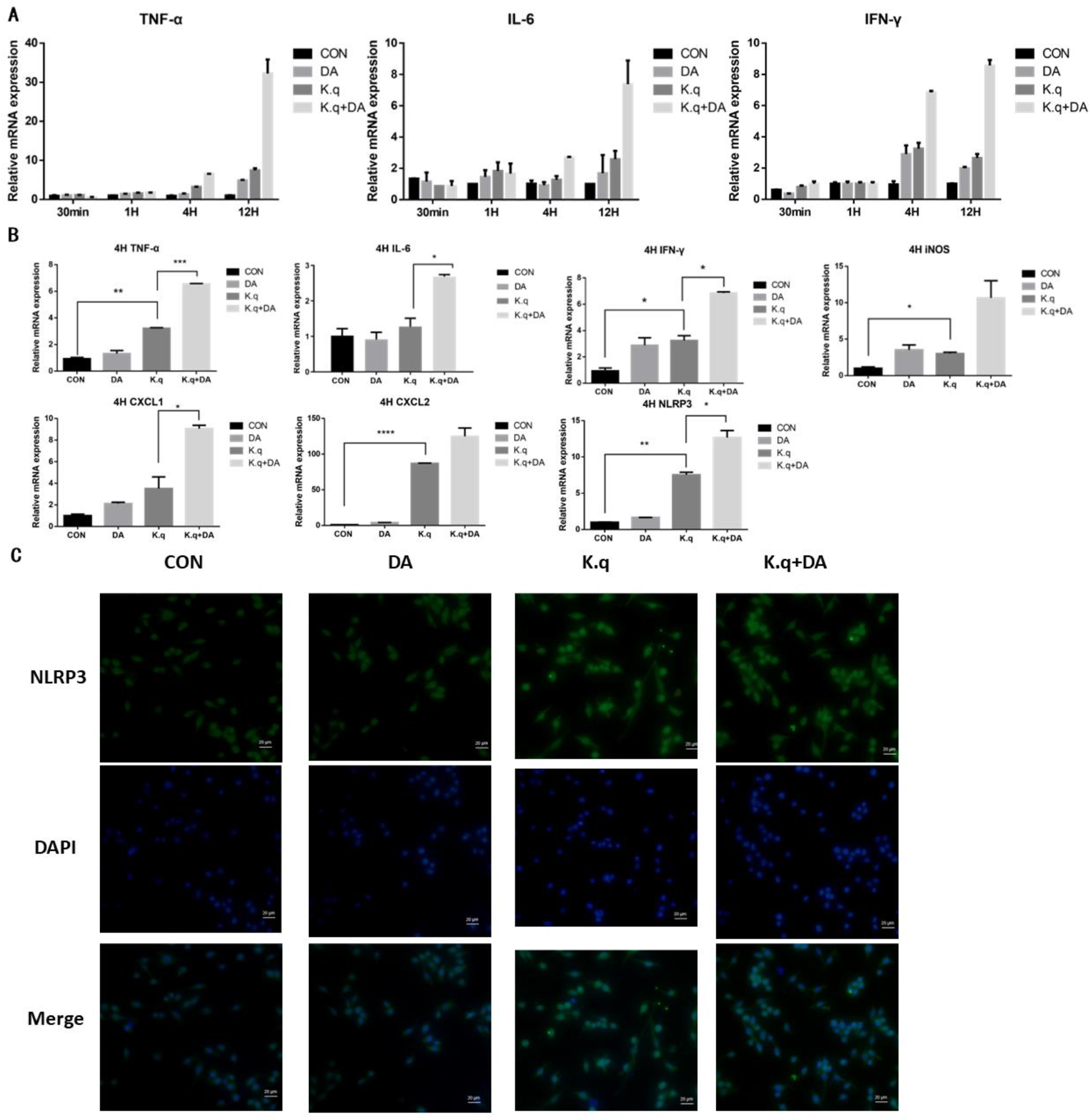
The detection of inflammatory factors in RAW 264.7. RAW264.7 were cultured in DMEM medium containing 10% FBS, in the absence (control guoup) or presence (dopamine group) of 500 μg/ml dopamine. RAW264.7 cells infected with *K. quasivariicola* at an MOI of 10:1 was considered as K. q group, and infected cells treated with 500 μg/ml dopamine was considered as K.q+DA group. The cytokines expressed in RAW264.7 cells were detected by qPCR (A,B) and immunofluorescence analysis (C) after 1, 4 and 12 hours, respectively.

### *K. quasivariicola* led to the lung infection and proinflammatory response in mice, and dopamine made outcome of *K. quasivariicola* infection worsen

C57/BL6 mice infected with *K. quasivariicola* were used to create acute lung injury models (Fig. 3A). After 48 hours-infection, pathological changes in the lung were observed. In comparison to the control group, lung tissue of mice infected with *K. quasivariicola* showed structural disruption and edema of alveolar epithelial cells, as well as neutrophil infiltration (Fig. 3B). It demonstrated that *K. quasivariicola* was a potential pathogen in pneumonia. It was noted that the inflammatory effects of *K. quasivariicola* were obviously aggravated when dopamine was administrated. The administration of dopamine in *K. quasivariicola* infected mice induced a complete destroyed alveolars, along with massive neutrophil infiltration and erythrocyte diapedesis (Fig. 3B).

**Figure 3:**
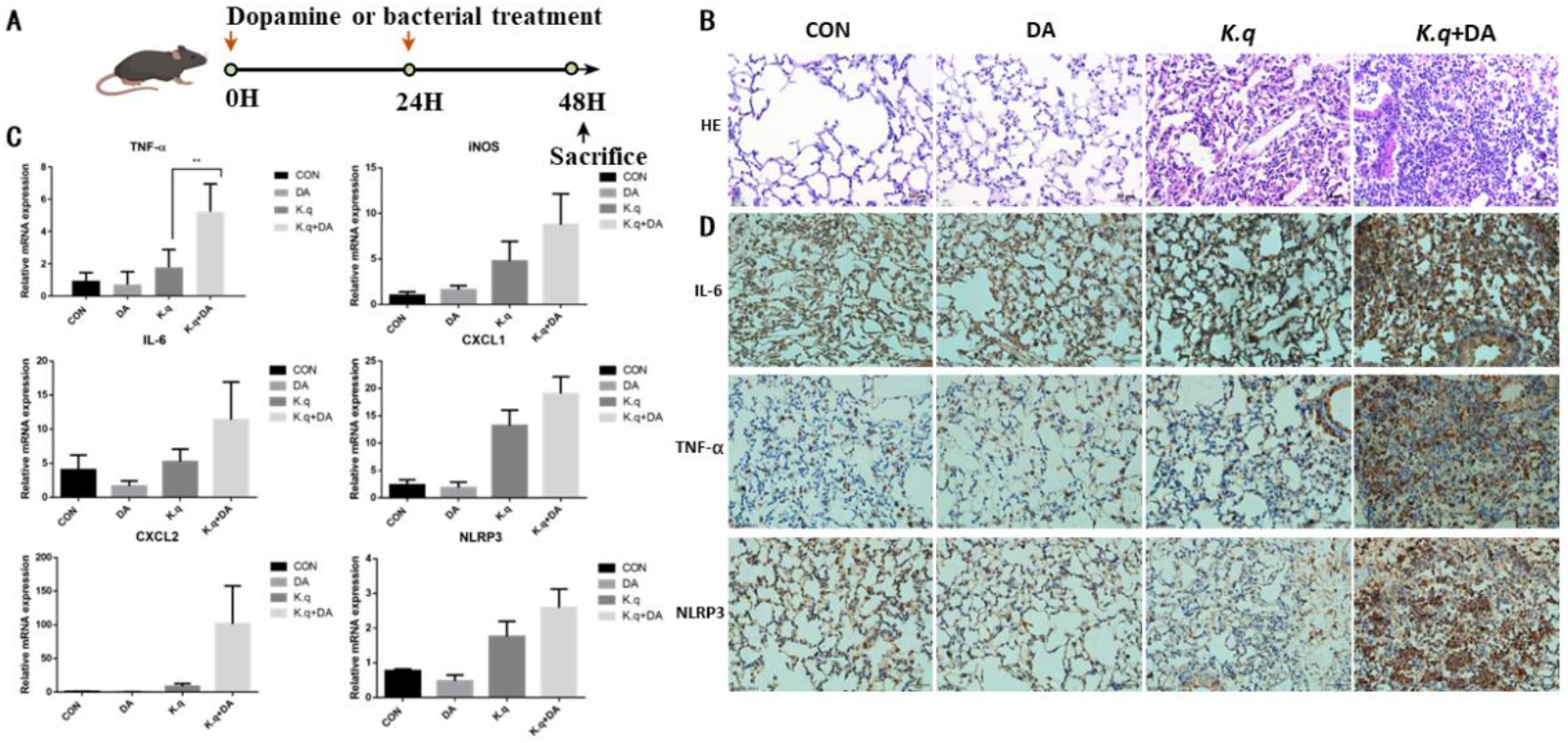
Lung histology of *K. quasivariicola* infected mice. Diagram of the Animal Experiment Plan. (B) The mice were randomly divided into four groups: (1) control group (CON), (2) dopamine group (DA), (3) *K. quasivariicola* group (K.q), and (4) dopamine + *K. quasivariicola* group (K.q+DA). The control group received a saline intraperitoneal injection. The dopamine group was induced by intraperitoneal injections of same amount of 50 μg/g of dopamine. In the *K. quasivariicola* group, 1×10^8^ CFU of *K. quasivariicola* in 50 μl saline was administrated by oropharyngeal instillation. And in the *K. quasivariicola* + dopamine group, in addition to *K. quasivariicola* infection, 50 μg/g of dopamine was also given to mice. For each group, a second administration was performed 24 hours later. After 48 hours of treatment, lung tissues from each group were collected and stained with H&E. (C-D) The detection of inflammatory factors in mice. The lung tissues of each group were collected after 48 hours of treatment. (C) Total RNA was isolated after homogenizing the lung tissues. The mRNA level of cytokines in mice lung were detected by qPCR. (D) The immunohistochemical assay of lung tissue with the primary antibodies of anti-IL-6, anti-TNF-α and anti-NLRP3.

The expression of proinflammatory factors in C57/BL6 mice infected with *K. quasivariicola* were also measured. Consistent with cell experiment, increased expression of IL-6, iNOS, TNF-α, CXCL1, CXCL2 and NLRP3 was observed in *K. quasivariicola* infected mice, and the expression level of these proinflammatory factors in infected mice increased significantly with dopamine treatment (Fig 3C). The immunohistochemical staining revealed that the expressions of IL-6 in lung tissue of *K. quasivariicola* infected mice was much higher than in the control group, and the expressions of IL-6 was further evaluated when dopamine was used (Fig 3D). The expression of TNF-α and NLRP3 inflammasome in lung tissue was also elevated during *K. quasivariicola* infection, but was mainly found in alveolar bronchioles (Fig 3D). However, dopamine administration resulted in a massive infiltration of immune cells with TNF- and NLRP3 expression in alveolar (Fig 3D).

## Discussions

### The crosstalk of dopamine and macrophage promoted the proliferation of K. quasivariicola

Dopamine was commonly used as a rescue drug in the Intensive Care Unit (ICU) to treat shock. Multiple studies have indicated that dopamine could regulate bacterial growth. *Cuvas Apan* et al. (21) found that dopamine at clinically used concentrations could decrease the growth of *Staphylococcus aureus, Staphylococcus epidermidis, Candida albicans, E. coli* and *P. aeruginosa*, and this effect was more prominent at higher concentrations. However, some other research showed dopamine promote the microbial growth of *S. Typhimurium, A. pleuropneumoniae* and *Yersinia ruckeri* (11, 22, 23). Our previous studies also proved dopamine can promote or inhibit the growth of a variety of bacteria derived from lung (Fig S1). It can be seen that the effects of dopamine on bacteria growth might vary between bacterial species.

However, the majority of these experiments were carried out in bacterial culture media, which ignored the effects of host immune systems. Therefore, in this study, the effects of dopamine on bacterial growth were investigated when *K. quasivariicola* co-cultured with macrophages. Our results showed dopamine inhibited the growth of *K. quasivariicola* in culture medium (Fig.1A,B), but significantly promoted the *K. quasivariicola* growth in the presence of macrophages (Fig.1C,D), indicating macrophage play important roles in the interactions between dopamine and *K. quasivariicola*.

The same phenomenon was also observed in *S. typhimurium*. Dichtl *et al*. (11) proved dopamine stimulated the intracellular growth of *S. Typhimurium* by increasing the bacterial iron acquisition. Iron was well known to be essential for bacterial growth and the limitation of iron could impair the replication of bacteria. One of the strategies for bacteria to collect iron was to synthesize endogenous siderophores, an iron-binding moieties, to capture and internalize ferric iron (24). It is interesting that a catechol core structure was found in siderophores. As the shared catechol structure, catecholamines were later be proved to transfer iron into bacteria, thereby promoted the bacterial growth in low-iron media (25). It was noting that dopamine could enhance the proliferation of bacteria only in iron-limited media, not in rich media (22).

Due to the vital of iron, host always to limit the availability of iron for microbial invaders during infection (26). Macrophages are major sites of iron storage in body, and maintain intracellular iron homeostasis by regulating the uptake, storage and release of iron. Studies showed dopamine stimulation could promote the iron accumulation in macrophages and results in the increase of intracellular bacterial growth (27). This explained why higher numbers of *K. quasivariicola* were found in macrophages in the presence of dopamine.

Moreover, the *K. quasivariicola* growth in the supernatant of RAW 264.7 cell culture medium with dopamine was also increased (Fig.1E), giving us a hint that there might be a kind of “growth stimulating factor” secreted from macrophages by the induction of dopamine. However, further researches are needed to confirm that.

### Dopamine triggered the intense inflammation response due to the facilitation of K. quasivariicola growth

To determine the pathogenic potential of *K. quasivariicola*, a mice pneumoniae model by *K. quasivariicola* infection was established. Our findings revealed that *K. quasivariicola* infection could trigger an inflammation response in lung, however, the administration of dopamine significantly aggravated the inflammation. In infected lung tissues without dopamine administration, although there were a number of infiltrating inflammatory cells, the expression of proinflammatory cytokine and NLRP3 were at lower level. However, once dopamine was added, large quantities of proinflammatory cytokines were produced, resulting in the recruitment of mass inflammatory cells and an intense inflammation.

Instead, previous studies have reported dopamine possessed anti-inflammatory effects. It was showed that dopamine could inhibit LPS-induced activation of NLRP3 inflammasome and reduce the subsequent production of caspase-1 and IL-1β (28). Further studies indicated dopamine negatively regulated the NLRP3 inflammasome through G protein pathway. The binding of dopamine and its receptor (D1-like receptor), a G protein-coupled receptor, could stimulate the activity of adenylate cyclase, and then promote the production of cAMP. cAMP can directly bind to NLRP3 to trigger the ubiquitination of NLRP3, thereby lead to an autophagy-mediated degradation of NLRP3 (29).

Interestingly, our results demonstrated that the administration of dopamine contributed in more severe inflammation, both *in vitro* and *in vivo*. The same phenomenon has been reported by Dicht *et al*. (11), which *S. typhimurium* infected mice receiving dopamine showed a significantly increased immune response. Due to the anti-inflammatory effects of dopamine, it can be thought that dopamine does not directly influence the immune response during *K. quasivariicola* infection. From our results, we considered that dopamine triggered the severe inflammatory response by significantly promoting the growth of *K. quasivariicola*, and the sharply increased bacterial load then triggering the release of proinflammatory cytokines in large amounts.

Dopamine is the first-line vasoactive drug used in antishock therapy for critically ill patients. Our research indicated that during *K. quasivariicola* infection, the administration of dopamine led to overproduction of proinflammatory cytokines and provoked the risk of cytokine storm, therefore, for *K. quasivariicola* infected patients, the administration of dopamine was inappropriate. Previous studies had reported that dopamine could enhance some certain bacterial growth in macrophage by increasing the iron acquisition. It was noting that Stefanie *et al*. (27) had reported that only dopamine, but not other catecholamines, could increase the intracellular iron content of macrophages. Therefore, we recommended that for critical patients with *K. quasivariicola* infection, epinephrine or norepinephrine should be first choice. Our studies could provide references for precision medication in critical patients, preventing the aggravating infections and reducing the mortality.

### An in vitro model of microbes-drugs-host immune cells was proposed for inhibitor screening

Of noting, dopamine showed contradiction effects on bacterial growth while *K. quasivariicola* mono-cultured in medium or co-cultured with macrophages, indicating macrophages must contributed to the interaction between dopamine and *K. quasivariicola*. This made us reflect the current studies of inhibitor screening. It should be noted that previous studies about the interactions between microbes and monomeric compounds were always limited in culture medium. Once the compound could inhibit the growth of bacteria in medium, it was considered as a candidate antibiotic. Nevertheless, there were more sophisticated communications between microbes and drugs in the niche of pathogens colonized in hosts. In the studies of inhibitor screening, unsatisfactory results or even opposite results would be achieved when host immune response were ignored. Therefore, an *in vitro* model which contains microbes-drugs-host immune cells, which could better mimic *in vivo* environment, was proposed for inhibitor screening (Fig 4). In the *in vitro* model of microbes-drugs-host immune cells, the growth of bacteria was more similar to that *in vivo*, and the changes of inflammatory factors caused by bacteria or drugs could be detected at the same time. Therefore, the development of *in vitro* model is of great importance for study the interactions between microbes and drugs.

**Figure 4:**
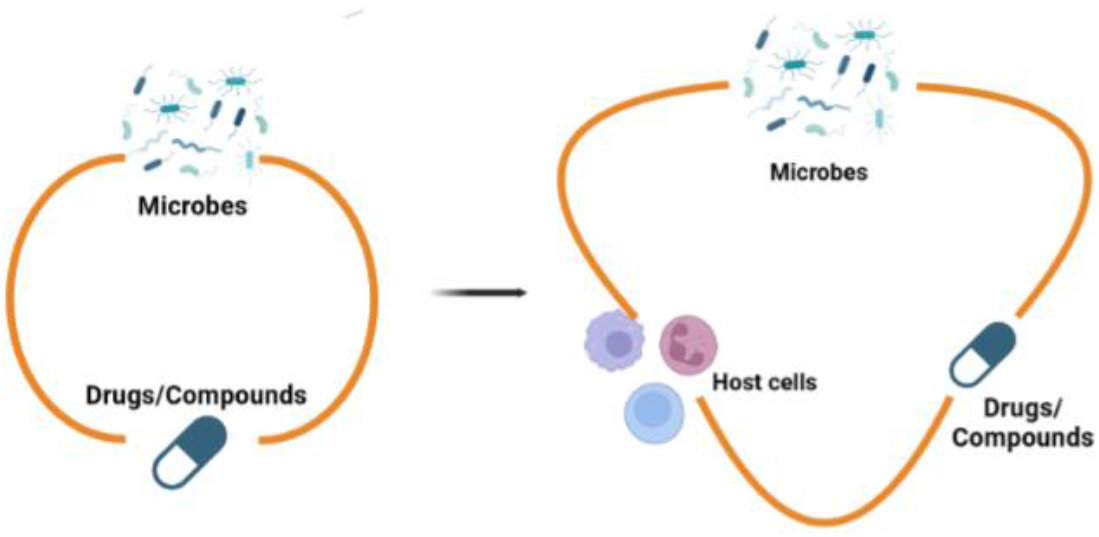
The *in vitro* model of microbes-drugs-host immune cells.

In conclusions, our researches demonstrated that *K. quasivariicola* was one of the potential pathogens for pneumonia and led to a pulmonary inflammation. The administration of dopamine could aggravate the inflammation reactions by promoting the proliferation of *K. quasivariicola* in the presence of macrophages. Here we made a recommendation that critically ill patients with *K. quasivariicola* infection should be treated with other catecholamines such us epinephrine or norepinephrine instead of dopamine. These may provide us with references for precision medication in critical patients. Furthermore, an *in vitro* model of microbes-drugs-host immune cells for inhibitor screening was proposed. The development of *in vitro* models that better mimic *in vivo* environments is of great importance for study the interactions between microbes and drugs.

## Materials and methods

### Bacterial strains and growth conditions

The *K. quasivariicola* strain used in this study was isolated from sputum sample of a pneumonia patient at ICU of Dalian Municipal Central Hospital, and identified by full-length sequencing of 16S rRNA gene (Fig S2). The bacteria were cultured in DMEM containing 10% FBS at 37°C. The medium was supplemented, as needed, with 500µg/ml dopamine. The study was approved by the Ethics Committee of the Dalian Central Hospital (Ethical approval number: 2017-030-01). Informed consent was obtained from the subject. The clinical samples were collected in accordance with the approved guidelines.

For the growth of *K. quasivariicola* in cell culture supernatants, the RAW264.7 cells (2×10^6^) were cultured in DMEM containing 10% FBS and 500 µg/ml dopamine at 37°C. The cell cultures without dopamine were served as control. After 4 hours-cultivation, the supernatants were harvested and sterilized by passing it through a 0.22 μm filter (Millipore). The *K. quasivariicola* strain was then cultivated in supernatants at 37°C for 8 hours before CFU counting was done.

### Bacterial growth with RAW264.7 macrophages

In 24-well plates, RAW264.7 cells (2×10^5^ per well) were seeded and cultured in DMEM containing 10% FBS and 500 ug/ml dopamine. The RAW264.7 control group was subjected to the same culture conditions as the experimental group, with the exception of the addition of dopamine. After culturing for ∼12 hours, RAW264.7 macrophages were infected with *K. quasivariicola* at a 10:1 MOI for 4 hours of incubation. The cell culture medium were then used to count the CFUs of *K. quasivariicola*. Meanwhile, RAW264.7 cells were collected and rinsed three times with PBS to remove extracellular bacteria. The cells were lysed with 500 μl of 0.03% SDS on ice. After homogenization, 10-fold serial dilutions were plated onto TSA plates to determine the CFU.

### RNA isolation and RT-qPCR

RAW264.7 cells (4×10^5^ per well) were cultured in 12-well plates in DMEM containing 10% FBS. After ∼12 hours, the cells were collected and resuspended in fresh DMEM medium containing 10% FBS. Then the cells were divided into 4 groups: RAW264.7 cells cultured in the absence of dopamine was considered as control group (CON), and the cells cultured with 500 μg/ml dopamine was considered as the dopamine group (DA). RAW264.7 cells infected with *K. quasivariicola* at an MOI of 10:1 was considered as K. q group (*K*.*q*), and the infected cells treated with 500 μg/ml dopamine was considered as Kq+DA group (*K*.*q*+DA). Cells were cultured for an additional 1, 4 or 12 hours, and then harvested for RNA isolation and RT-qPCR detection.

Total RNA of RAW264.7 cells were isolated using the RNAiso plus (TaKaRa, Japan) according to the manufacturer’s instructions. The isolated RNA (1 μg) was immediately reverse transcribed into cDNA using the AG Evo M-MLV RT Kit with gDNA Clean for qPCR Kit (Accurate Biotechnology, China). The expressions of TNF-α, IL-6, CXCL1, CXCL2, IFN-γ, β-actin, IL-8, iNOS, IL-17, IL-18 were analyzed by RT-qPCR using the specific primers list in table 1. Data from three independent experiments were used for statistical analysis.

**Table 1:**
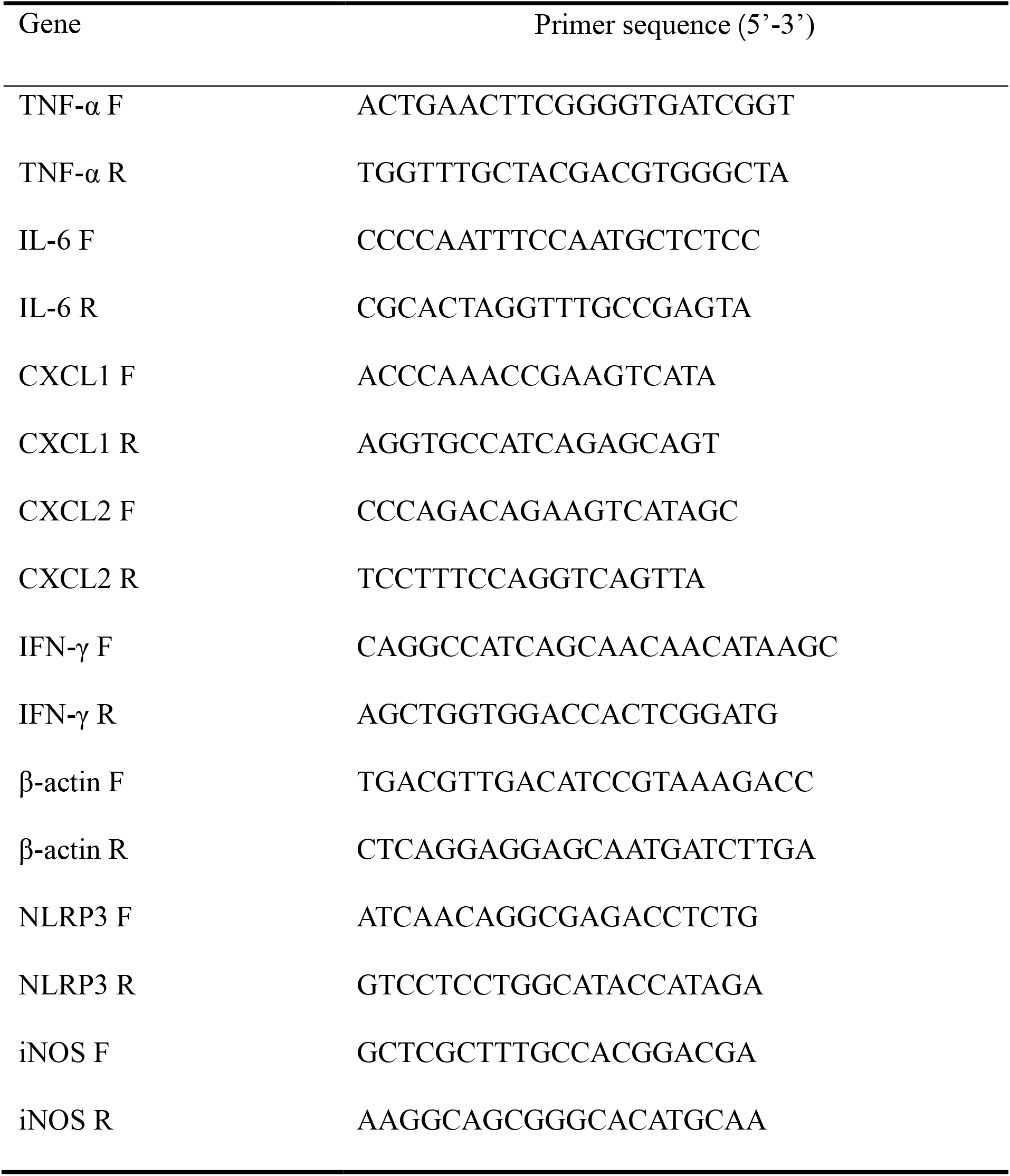
Primers used for qPCR.

### Immunofluorescence Staining

Cells were fixed in 4% paraformaldehyde for 15 minutes at room temperature before being permeabilized in 0.1% Triton X-100 for 15 minutes. After rinsing with PBS, cells were blocked with 1% BSA for 60 min at room temperature. After blocking, cells were incubated with anti-NLRP3 antibody (boster, Wuhan, China) overnight at 4°C. Cells were then washed with PBS to remove the excessive antibodies, and incubated with fluorescent secondary antibodies for 1 hour at 37°C. Samples were counterstained with DAPI (Invitrogen, Carlsbad, CA, United States) for 15 minutes to visualize the nuclei. Finally, the images were analyzed using an inverted fluorescent microscope (Olympus, Japan).

### Mouse model of acute lung injury

All animal experiments were approved by the Committee on the Ethics of Animal Experiments of Dalian Medical University (Permission number: SYXK (Liao) 2018-0007) and were performed in strict accordance with the recommendations. Sixteen C57/BL6 mice (aged 8 weeks, weight 20-25 g) were randomly divided into four experimental groups: (1) control group (CON), (2) dopamine group (DA), (3) *K. quasivariicola* group (k.q), and (4) *K. quasivariicola* + dopamine group (k.q+DA). In control group, the mice were intraperitoneal injection with saline. In dopamine group, mice were induced by intraperitoneal injections of same amount of 50 μg/g of dopamine. In *K. q* group, 1×10^8^ CFU of *K. quasivariicola* in 50 μl saline was administrated by oropharyngeal instillation. And in *K. quasivariicola* + dopamine group, in addition to *K. quasivariicola* infection, 50 μg/g of dopamine was also given to mice. For each group, a second administration were performed 24 hours later. All the mice were ready for the sacrifice after 48 hours of first treatment (Fig 3A). The homogenates of lung tissue were used for RNA isolation and the expressions of TNF-α, IL-6, CXCL1, CXCL2, IFN-γ, β-actin, IL-8, iNOS, IL-17, IL-18 were analyzed by RT-qPCR as described previously.

### Lung histopathology

The lung tissue was fixed in 4% paraformaldehyde, paraffin-embedded, and cut into slices for routine HE staining and immunohistochemical assays. Briefly, paraffin-embedded tissues were sliced into 4 μm thick slices, dewaxed and gradually rehydrated with ethanol. The sections were then treated for 20 minutes at 95 °C with DAKO Target Retrieval solution (Dako) for epitope retrieval. Following that, the sections were treated with 0.3% hydrogen peroxide and probed overnight with anti-IL-6, anti-TNF-α and anti-NLRP3 primary antibodies. The secondary antibodies were then added and incubated for 30 minutes. The proteins were detected and visualized with streptavidin-HRP conjugates and DAB substrate solution. For HE staining, the deparaffinized sections were stained with hematoxylin, then treated with 1% acid alcohol, and finally with 1% eosin.

## Acknowledgments

This work was supported by grants from the National Natural Science Foundation of China (81902037). The funders had no role in study design, data collection and interpretation, or the decision to submit the work for publication.

The authors declare that there is no conflict of interests.

QY, JK, YM and TM contributed to conception and design of the study. JK and QY wrote the manuscript. QY and YM leaded the Writing-Review & Editing. XL, LC and XL collected the samples and performed the experiments. XW, RL and SF performed the data analysis. All authors contributed to the article and approved the submitted version.

## Supplementary data

**Supplementary Figure 1:**
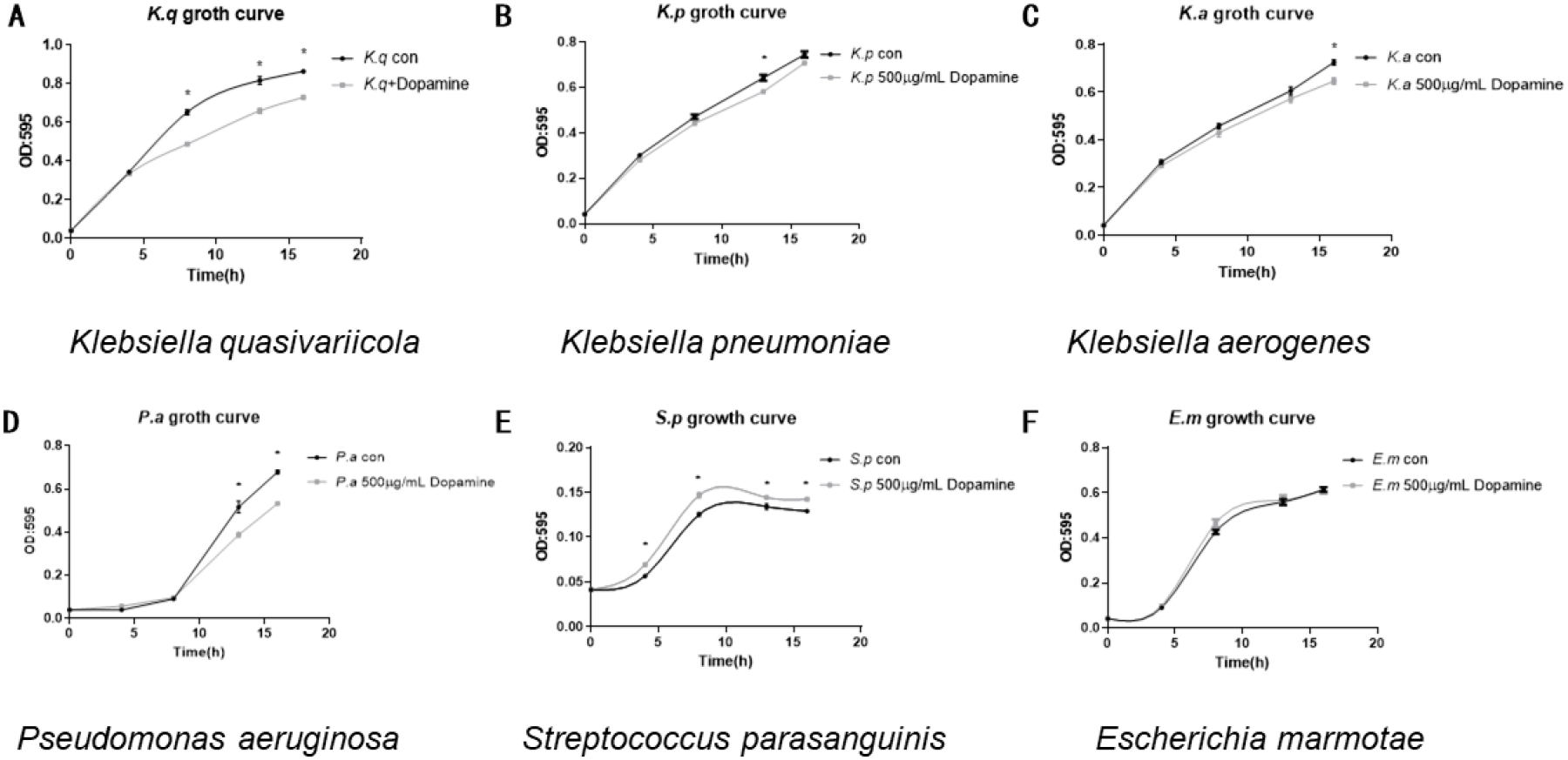
The growth curve of different strains in TSA medium. The different strains were cultured in TSA medium. Black, bacteria cultured only; Gray, bacteria cultured with 500 µg/ml dopamine. The OD620 of cultures were monitored every 4 hours. Each sample was assayed in triplicate. Asterisk (*) means the difference between two groups was statistically significant. Dopamine inhibited the growth of *Klebsiella quasivariicola* (A), *Klebsiella pneumoniae* (B), *Klebsiella aerogenes* (C) and *Pseudomonas aeruginosa* (D), however, promoted the growth of *Streptococcus parasanguinis* (E) and *Escherichia marmotae* (F).

**Supplementary Figure 2:**
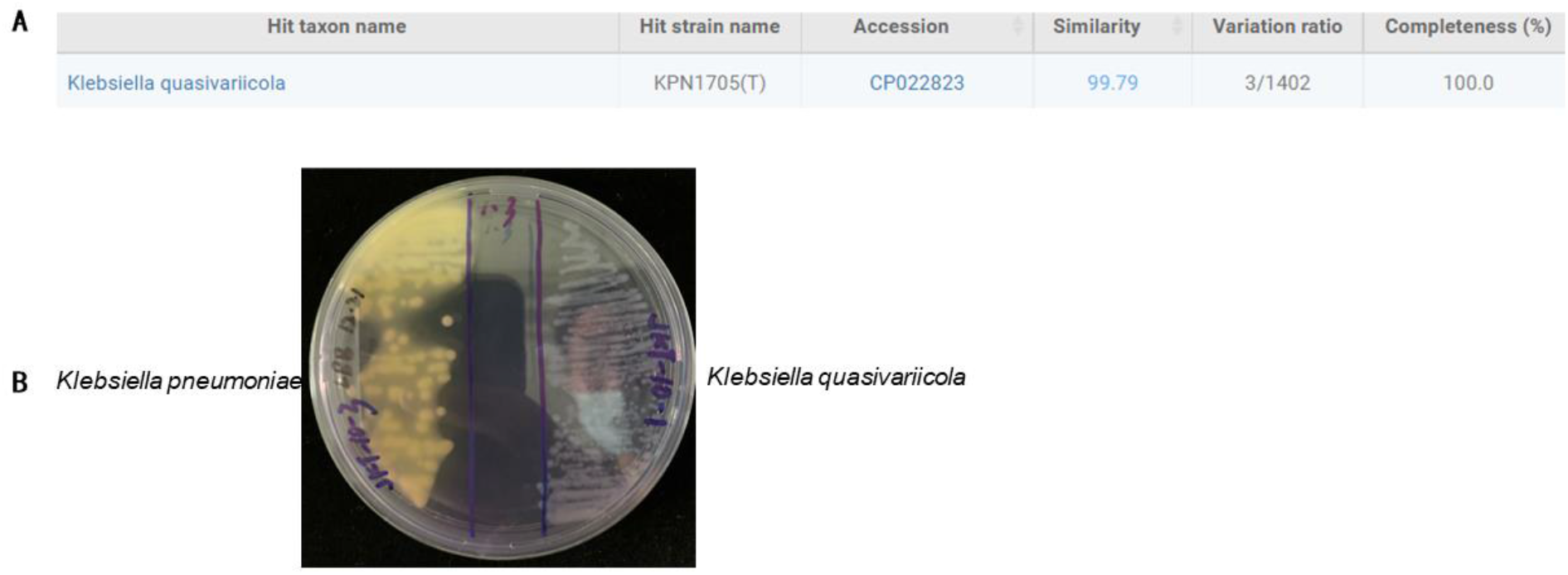
The 16S rRNA gene search in EzBioCloud Database and Colony morphology of *K. quasivariicola*. (A) 16S rRNA sequencing of *K. quasivariicola*. (B) Colony morphology of *K. quasivariicola* (right) and (left) on LBB plate. There was the pH indicator of bromocresol violet on LBB plate, which could change the color from purple to yellow under acidic conditions. *K. quasivariicola* and *K. pneumoniae* showed different morphology on LBB, indicated the acid producing capacities of these two strains were different.

