## Supplementary figures and images for "Dopamine promotes *Klebsiella quasivariicola* proliferation and inflammatory response in the presence of macrophages"

### Supplemental Figure

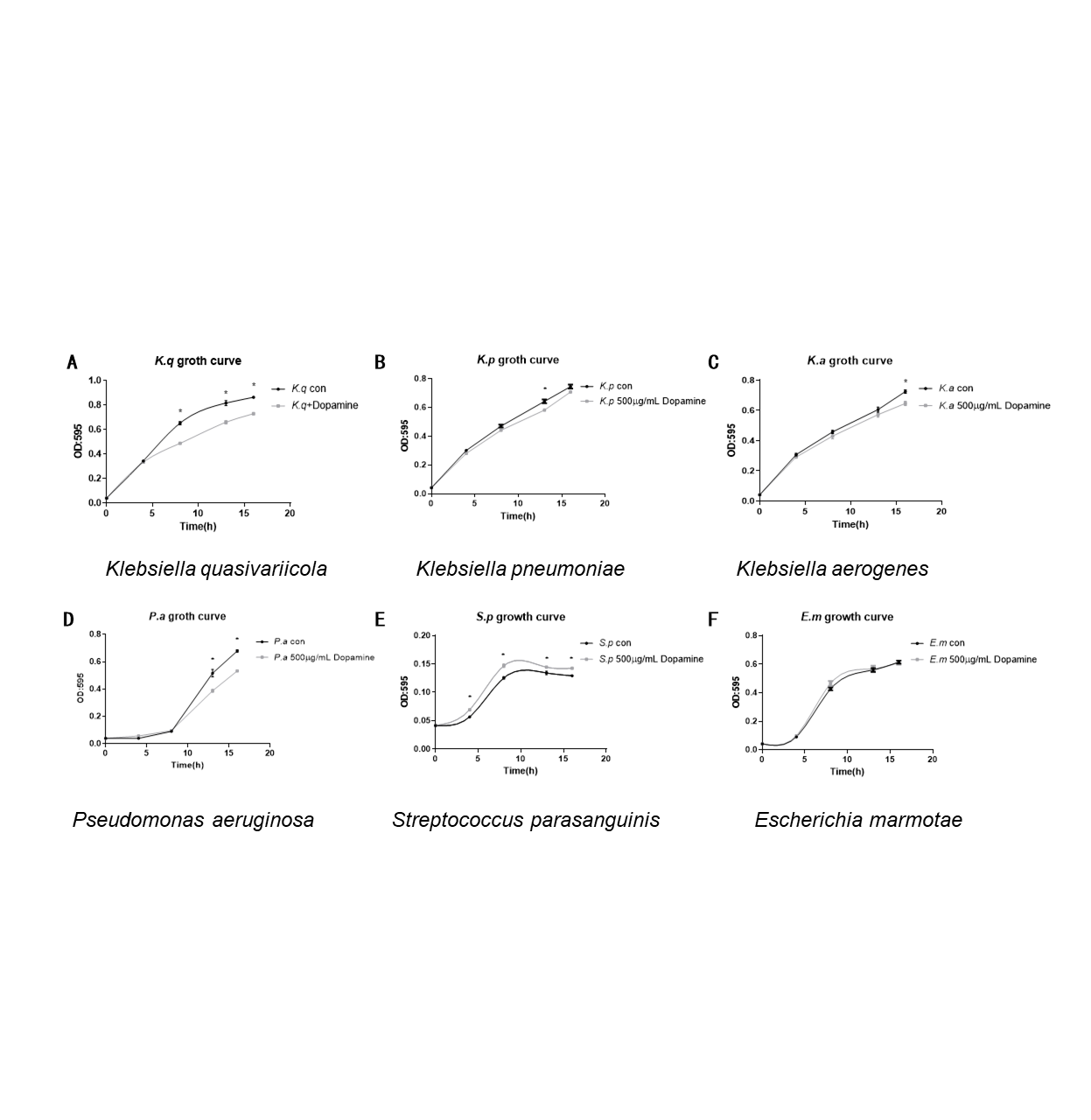

### Supplemental Table

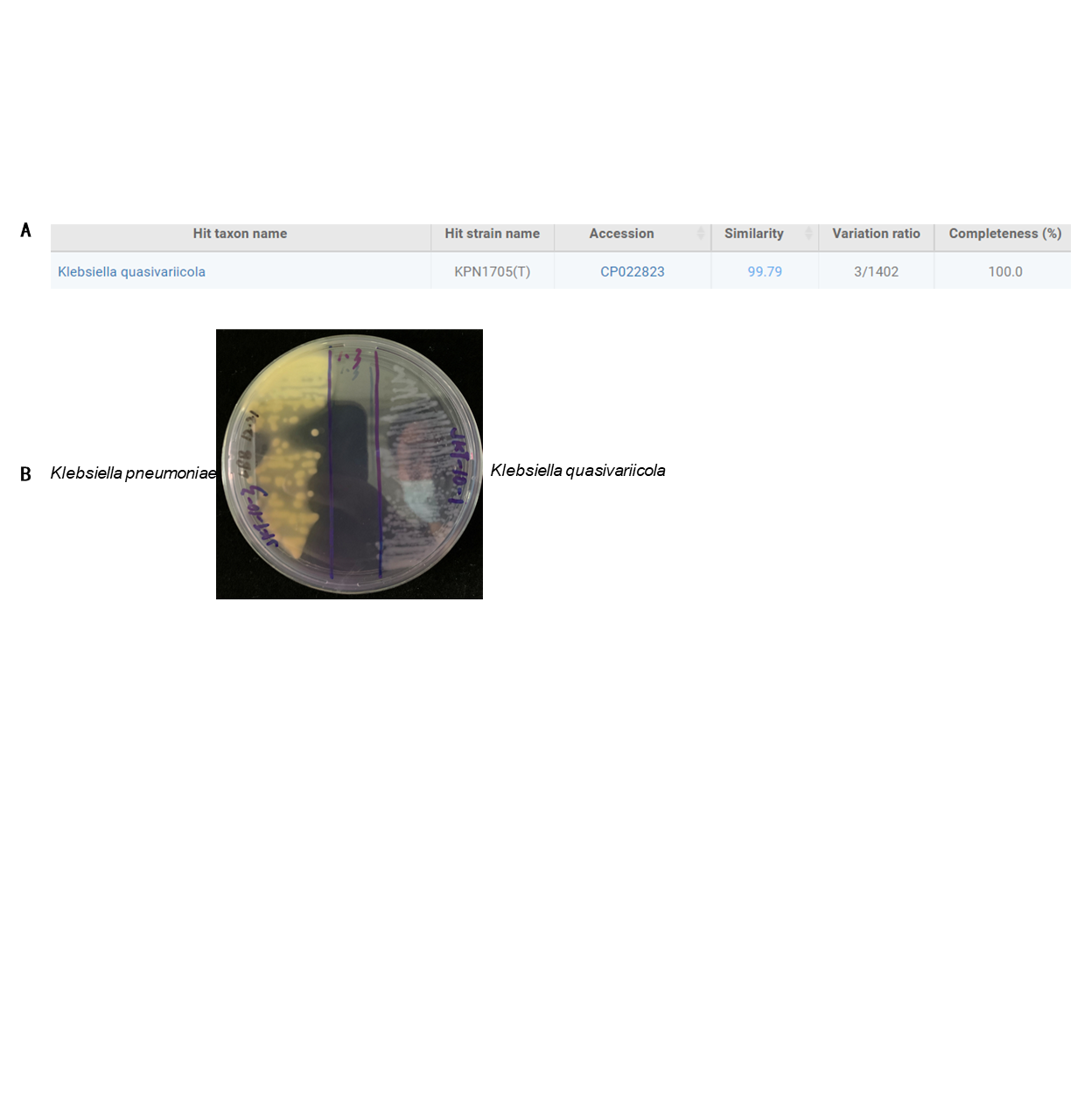
